# TGF-β1 inhibits cholesterol metabolism in hepatocytes to facilitate cell death, EMT and signals for HSC activation

**DOI:** 10.1101/2023.08.14.552900

**Authors:** Sai Wang, Frederik Link, Mei Han, Roman Liebe, Ye Yao, Seddik Hammad, Anne Dropmann, Roohi Chaudhary, Anastasia Asimakopoulos, Marinela Krizanac, Ralf Weiskirchen, Yoav I. Henis, Marcelo Ehrlich, Matthias P. A. Ebert, Steven Dooley

## Abstract

**Objective:** Transforming growth factor-β1 (TGF-β1) plays important roles in chronic liver diseases, including metabolic dysfunction-associated steatotic liver disease (MASLD), which involves various biological processes including dysfunctional cholesterol metabolism contributing to progression to metabolic dysfunction-associated steatohepatitis (MASH) and hepatocellular carcinoma (HCC). However, how TGF-β1 signaling and cholesterol metabolism affects each other in MASLD is yet unknown.

**Design:** Changes in transcription of genes associated with cholesterol metabolism were assessed by RNA-Seq of AML12 cells and mouse primary hepatocytes (MPH) treated with TGF-β1. Functional assays were performed on AML12 cells (untreated, TGF-β1 treated, or subjected to cholesterol enrichment (CE) or depletion (CD)), and on mice injected with adeno-associated virus 8 (AAV8)-Control/TGF-β1.

**Results:** TGF-β1 inhibited mRNA expression of several cholesterol metabolism regulatory genes, including rate-limiting enzymes of cholesterol biosynthesis in AML12 cells, MPHs, and AAV8-TGF-β1-treated mice. Total cholesterol levels in AML12 cells, as well as lipid droplet accumulation in AML12 cells and AAV-treated mice were also reduced. Smad2/3 phosphorylation following 2 h TGF-β1 treatment persisted after CE or CD and was mildly increased following CD, while TGF-β1-mediated AKT phosphorylation (30 min) was inhibited by CE. Furthermore, CE protected AML12 cells from several effects mediated by 72 h incubation with TGF-β1, including EMT, actin polymerization, and apoptosis. CD mimicked the outcome of long term TGF-β1 administration, an effect that was blocked by an inhibitor of the type I TGF-β receptor. Additionally, the supernatant of CE- or CD-treated AML12 cells inhibited or promoted, respectively, the activation of LX-2 hepatic stellate cells.

**Conclusion:** TGF-β1 inhibits cholesterol metabolism while cholesterol attenuates TGF-β1 downstream effects in hepatocytes.

## Introduction

The multifactorial metabolic dysfunction-associated steatotic liver disease (MASLD) is the most common cause of chronic liver disease ^1^. It encompasses a spectrum of pathological conditions ranging from simple steatosis, metabolic dysfunction-associated steatohepatitis (MASH) and fibrosis/cirrhosis, which can further progress to hepatocellular carcinoma and liver failure. At the early stage of MASLD development, hepatocytes accumulate various lipids (triglycerides, free fatty acids, and cholesterol). This induces steatosis and increases the vulnerability to injury caused by lipotoxicity, mitochondrial dysfunction, cellular stress and inflammation, thought to be required to progress to MASH ^2^. In this process, the defective hepatocyte regeneration for the replenishment of dead cells and the adverse factors, including inflammatory factors, dietary factors, and lipopolysaccharide (LPS) are the proposed hits that drive the progression of MASLD to fibrosis ^3^. Later on, these insults lead to the activation of hepatic stellate cells (HSCs), accumulating collagen and extracellular matrix (ECM) production/deposition and portal hypertension, thus worsening liver fibrosis to cirrhosis and even HCC ^4^.

Hepatic lipotoxicity implies exposure to, or accumulation of, certain lipid species within hepatic cells that may initiate MASLD ^5^. Potentially lipotoxic molecules include cholesterol ^6, 7^, free fatty acids ^5^, and diacylglycerol ^8^. 50% of total cholesterol biosynthesis in humans occurs in the liver, where dysregulation of cholesterol homeostasis in MASLD leads to cholesterol accumulation ^9^. Excess dietary cholesterol has been shown consistently to cause the development of experimental MASH in different animal models, including high-fat diet, obesity, and hyperphagia ^10–12^. Additionally, cholesterol is essential for membrane structure and function, as it packs with phospholipids to form lipid microdomains, affects membrane fluidity and modulates membrane trafficking, host-pathogen interaction, and signal transduction ^13^, including transforming growth factor-β (TGF-β) signaling ^14–18^. TGF-β1 is a 25-kDa homodimeric peptide, playing significant roles in various biological processes, such as extracellular matrix formation, EMT, cell proliferation and differentiation, apoptosis, immune and inflammatory responses, and metabolic pathways ^19–23^. TGF-βs signal through binding to plasma membrane complexes of type II (TβR-II) and type I (TβR-I) TGF-β receptors ^24–26^. Apart of signaling to the canonical Smad2/3 pathway ^25, 27, 28^, and depending on the cell type and context, TGF-β also stimulates several non-Smad pathways ^29–33^, of which a prominent one in hepatocytes is the phosphoinositide 3-kinase (PI3K)/protein kinase B (AKT) pathway ^17^. Statin treatment is associated with significant protection from steatosis, inflammation, and fibrosis among patients with MASLD and/or MASH ^34^. Moreover, diet with excess cholesterol mitigates liver fibrosis through oxidative stress-induced HSC-specific apoptosis in mice ^35^. Cholesterol-lowering statins were also reported to promote basal pancreatic ductal adenocarcinoma (PDAC) through activation of TGF-β signaling, leading to EMT ^36^. However, how cholesterol affects TGF-β signaling in the progression of MASLD is yet unknown, and it is unclear whether and how TGF-β affects cholesterol regulation.

In this study, we investigated the crosstalk between TGF-β1 signaling and cholesterol levels by using RNA-Seq analysis and functional assays in hepatocytes, hepatic stellate cells and WT mice overexpressing TGF-β1.

## Results

### TGF-β1 inhibits cholesterol metabolic processes according to RNA-Seq data analysis

In MASLD, more cholesterol accumulates in the liver and the homeostasis of its metabolism becomes dysregulated in hepatocytes ^37^. As TGF-β1 signaling is activated under these conditions ^38^, we investigated the effects of TGF-β1 on cholesterol metabolism in hepatocytes. Three control and TGF-β1 treatment groups of AML12 and MPH cells were subjected to RNA-Seq, and the genes involved in cholesterol metabolism were analyzed. TGF-β1 treatment for 24 h significantly inhibited transcription of multiple cholesterol metabolism genes, as shown in the heatmaps of AML12 and MPH cells (Figure 1A). These include, among others, *Apoa2*, *Apoc3*, *Cyp27a1*, *Fgl1*, *Insig2*, *Lcat*, *Hmgcs2*, *Aqle*, *Abcg5*, *Lss*, and *Cyp7a1*. The chord diagram demonstrates that these downregulated genes are related to cholesterol homeostasis, biosynthesis, metabolic processes, efflux, import, cholesterol binding, lipid transport, and lipoprotein metabolism (Figure 1B), hence supporting the critical role of TGF-β1 in reducing cholesterol metabolism in hepatocytes. Furthermore, networking analysis shows that the majority of these genes display co-expression and co-localization (Figure 1C). Analysis of the signaling pathways shows that there is also notable downregulation of genes involved in enhancing cholesterol esterification, bile acid signaling pathways, SREBP signaling pathway, and cellular response to cholesterol (Figure 1D, E).

**Figure 1.**
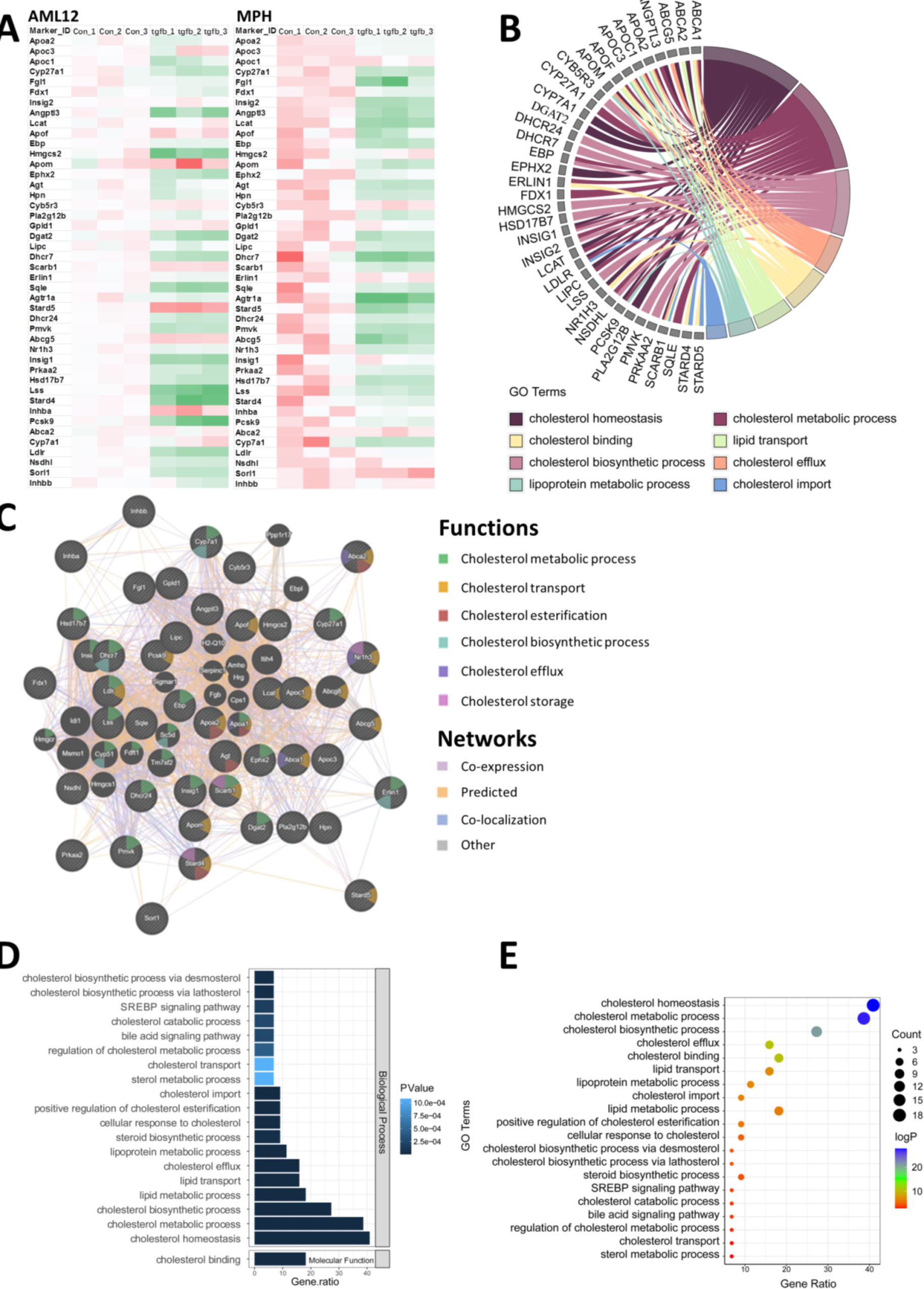
TGF-β1 inhibits cholesterol metabolism in hepatocytes. (A) Heatmap of the downregulated genes involved in cholesterol metabolism upon 24h TGF-β1 treatment in AML12 cells and MPHs. (number of independent experiments (n) = 3). (B) The chord diagram illustrates the main biological processes of the decreased genes. (C) Networking interpretation of the downregulated genes. (D, E) Signaling pathway analysis of the inhibited cholesterol metabolic genes.

### TGF-β1 inhibits cholesterol biosynthesis and accumulation in hepatocytes

According to the RNA-Seq analysis, TGF-β1 inhibits expression of multiple genes involved in cholesterol biosynthesis. This includes the 3 crucial players of the cholesterol biosynthetic pathway, Hmgcr, Sqle, and Lss, which were verified by RT-qPCR in AML12 cells (control vs. TGF-β1-treated). This result demonstrates that 24 h incubation with TGF-β1 had an inhibitory effect on mRNA expression of *Hmgcr* and *Sqle* compared to the control group (Figure 2A). The cholesterol assay shows a reduced concentration of cholesterol in AML12 cells incubated with 5 ng/ml TGF-β1 for 72 h, but not with 2 ng/ml TGF-β1 (Figure 2B). Treatment with 5 ng/ml TGF-β1 (72 h) also reduced lipid droplet accumulation in AML12 cells, identified by BODIPY staining (Figure 2C). Next, we examined the effects of overexpressing TGF-β1 by injecting mice with AAV8-TGF-β1 (or AAV8-Control) on cholesterol biosynthesis in the liver. RT-qPCR shows that overexpression of TGF-β1 reduced *Hmgcr*, *Sqle*, and *Lss* on the mRNA level (Figure 2D). In addition, lipid droplet accumulation was notably decreased in the liver tissue of AAV8-TGF-β1-injected mice compared to the control group (Figure 2E).

**Figure 2.**
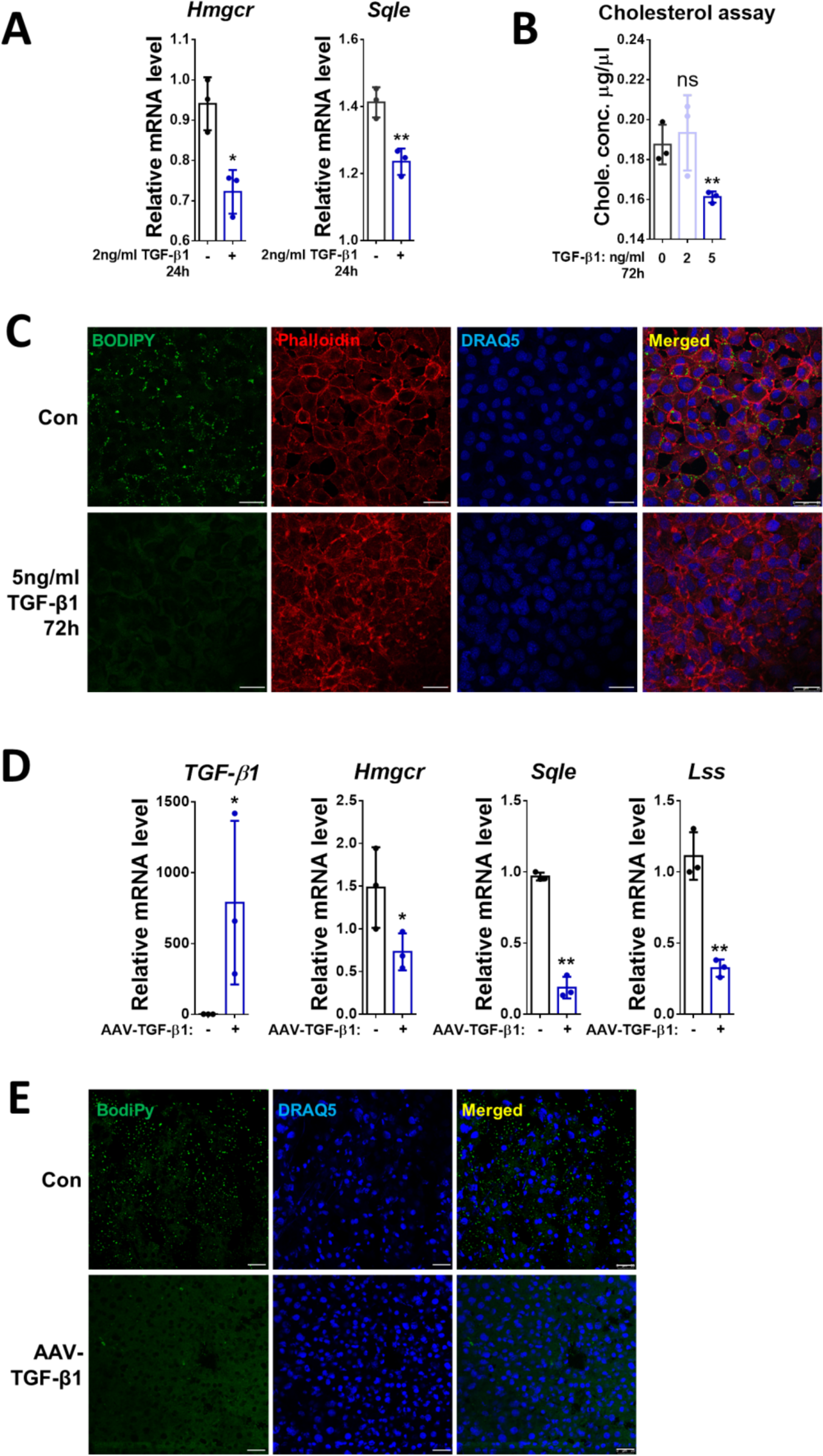
TGF-β1 inhibits cholesterol biosynthesis and accumulation in hepatocytes. (A) Relative mRNA levels of *Hmgcr* and *Sqle* were determined by RT-qPCR in AML12 cells treated with/without TGF-β1 for 24 h. (B) Total cholesterol concentration was examined in AML12 cells treated (or not) with 2 ng/ml or 5 ng/ml TGF-β1 for 72h. (C) Lipid droplet accumulation in AML12 cells was examined by BODIPY staining. Phalloidin was used for F-actin staining. (D) Relative mRNA levels of *TGF-β1*, *Hmgcr*, *Sqle, and Lss* were determined by RT-qPCR in liver tissue of mice treated with AAV8-control or AAV8-TGF-β1. (E) Lipid droplet accumulation in liver tissue of mice treated with AAV8-Control or AAV8-TGF-β1 was examined by BODIPY staining. For RT-qPCR, mouse *Ppia* was used as endogenous control. Bars represent mean ± SD (n=3). \**p*<0.05; \*\**p*<0.01, ns = p>0.05. For IF, DRAQ5 was used to stain the nuclei. Scale bars, 25 µm. Images were chosen representatively from 3 independent experiments.

### Cholesterol depletion and enrichment have different effects on TGF-β1-mediated Smad2/3 and AKT phosphorylation

Since TGF-β1 inhibits cholesterol biosynthesis (Figures 1 and 2), we next tested the effects of cholesterol on TGF-β1 signaling. Incubation with cholesterol-MβCD complex (5 mM MβCD, 300 µg/ml cholesterol) or with 50 µM lovastatin plus 50 µM mevalonate (MVL) induced CE or CD, respectively, in AML12 cells, compared to the control groups (Figure 3A). To investigate the effect of CE and CD on TGF-β1 signaling, AML12 cells were subjected to CE or CD treatments (or untreated; control) for 14-16 h as described under Materials and Methods. Subsequently, the cells were serum starved (0.5% FBS or LPDS) for 4 h, before treatment with or without 2 ng/ml TGF-β1 for 2 h (Smad2/3 signaling) or 30 min (AKT signaling). Western blot analysis for p-Smad2/3 and Smad2/3 showed that Smad2/3 phosphorylation in response to TGF-β1 (2 h) proceeded following either CE or CD treatments, with no effects for CE and some enhancement of p-Smad2/3 by CD (Figure 3B). On the other hand, CE treatment (but not CD) prevented TGF-β1-mediated (30 min) AKT phosphorylation at Ser473 (Figure 3C).

**Figure 3.**
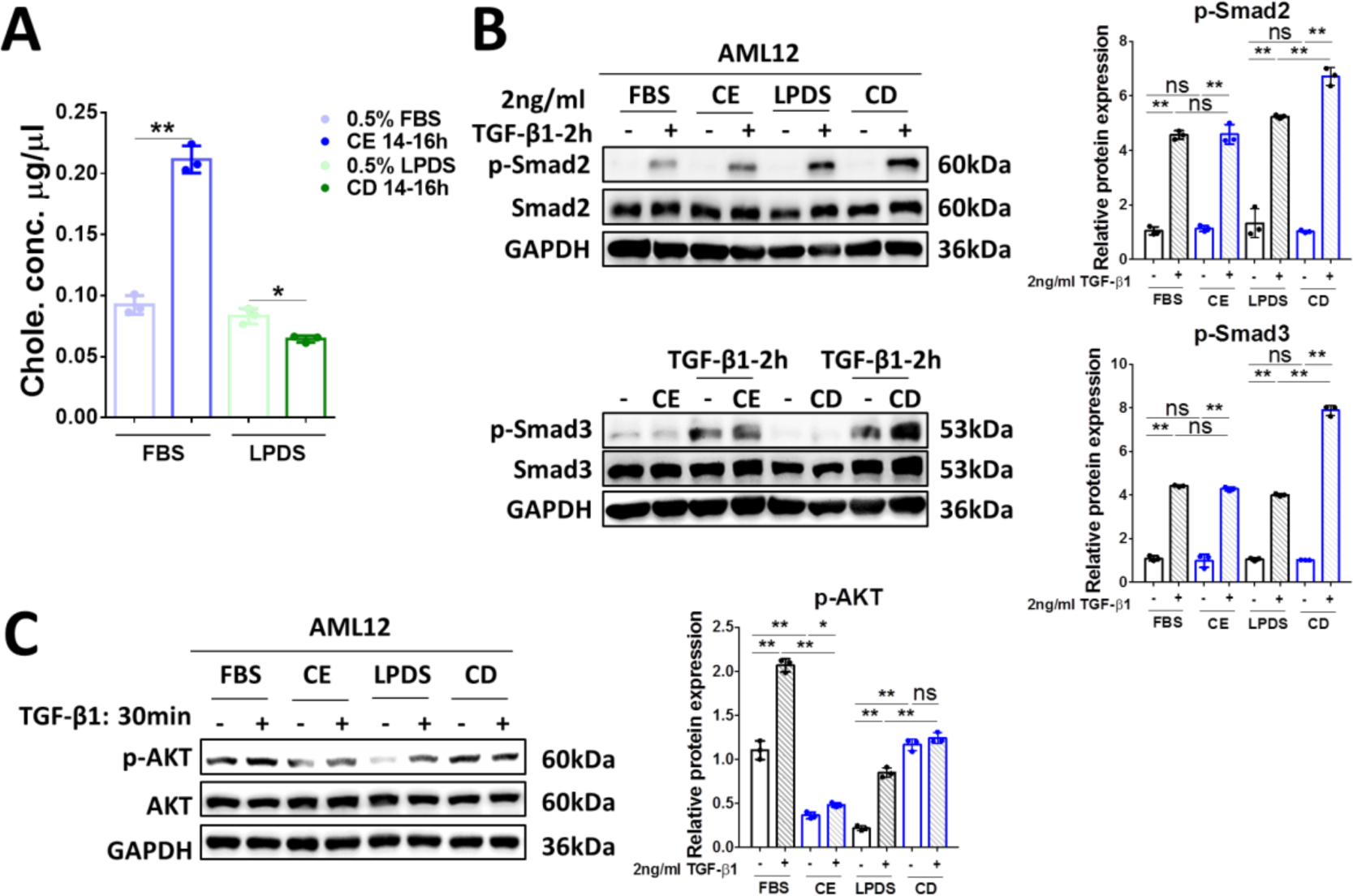
Alteration of the cholesterol level in AML12 cells affect differently TGF-β1-mediated signaling to Smad2/3 and to AKT. (A) Total cholesterol concentration was examined in AML12 cells treated with 5 mM cholesterol-MβCD complex (CE) or 50 µM lovastatin and 50 µM MVL (CD), respectively, for 14-16 h. (B) AML12 cells treated for CE, CD or untreated (control) were then stimulated with 2 ng/ml TGF-β1 for 2 h. The levels of p-Smad2 and total Smad2 (upper panels) or p-Smad3 and total Smad3 (bottom panels) were determined by Western blot analysis. (C) Cells treated to alter their cholesterol levels as above were incubated with 2 ng/ml TGF-β1 for 30 min, and subjected to Western blotting of p-AKT and total AKT. For RT-qPCR, mouse *Ppia* was used as endogenous control. Bars represent mean ± SD (n=3). \**p*<0.05; \*\**p*<0.01, ns = p>0.05. For Western blotting, GAPDH was used as a loading control. Quantification of protein expression was performed using Image J (National Institute of Health, Bethesda, MD).

### TGF-β1-induced EMT and stress fiber formation are inhibited by cholesterol enrichment and promoted by cholesterol depletion

AML12 cells undergo morphological changes following CE or CD treatments, which also affect morphological changes induced by prolonged incubation (72 h) with TGF-β1 (Figure 4A). We therefore investigated the effects of TGF-β1 on EMT and actin polymerization in AML12 (untreated, or subjected to CE or CE). TGF-β1 treatment for 72 h induced delocalization of E-Cadherin from the cell membrane, and its membrane localization was re-established by CE treatment (Figure 4B). However, E-Cadherin membrane localization was further decreased by CD treatment alone (Figure 4B). TGF-β1-induced fibronectin expression in the membrane was abrogated by CE, but further enhanced by CD, as shown by immunofluorescence (Figure 4C). RT-qPCR confirmed the upregulation of the mRNA levels of EMT markers *Twist1, Fn1*, and the downregulation of *Cdh1* by TGF-β1, and showed that they can be prevented by CE; on the other hand, they could be further phenocopied by CD (Figure 4D). E-cadherin expression at the protein level was decreased in the CD incubation group, as identified by Western blotting (Figure 4E). Although TGF-β1 inhibited *Cdh1* mRNA expression (Figure 4D, third histogram), there was no obvious decrease in its protein level (Figure 4E). According to the immunofluorescence results (Figure 4B), TGF-β1 mainly disrupted E-cadherin membrane localization rather than its expression. In addition, actin polymerization induced by TGF-β1 was reduced to normal levels by CE, while CD dramatically promoted this outcome, as indicated by staining of F-actin with phalloidin (Figure 4F). These findings indicate that CE prevented TGF-β1-induced EMT and actin polymerization, perhaps due to its inhibitory effect on AKT signaling (Figure 3C), which alters the balance between the stimulation of the Smad2/3 and AKT pathways. This did not occur in the case of CD treatment, where AKT phosphorylation was not inhibited while that of Smad2 may even be increased (Figure 3B). This different balance between the two pathways may be involved in the further aggravation by CD of TGF-β1-mediated outcomes in AML12 cells.

**Figure 4.**
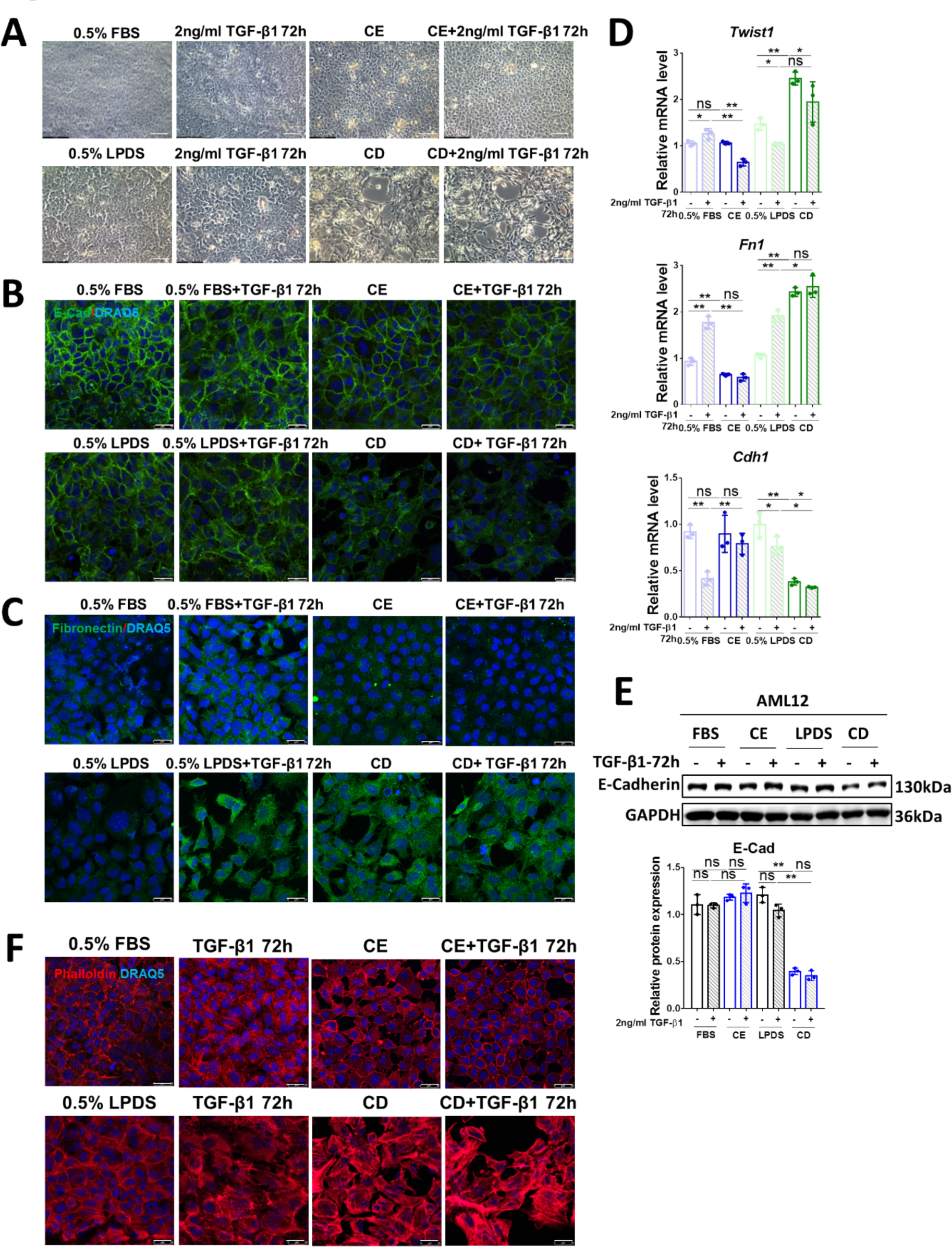
CE inhibits TGF-β1-induced EMT and stress fiber formation while CD promotes these effects. (A) Brightfield images of the morphology of AML12 cells treated with CE/CD combined with/without 2 ng/ml TGF-β1 for 72h. Scale bars, 50 µm. (B, C) The expression and location of E-Cadherin and Fibronectin were determined by immunofluorescence in AML12 cells treated with CE/CD combined with/without 2 ng/ml TGF-β1 for 72 h. (D, E) Relative mRNA levels of *Twist*, *Fn1*, and *Cdh1* (D) and protein levels of E-cadherin (E) were determined by RT-qPCR and Western blotting, respectively, in AML12 cells with the conditioned treatment as shown in the figure. (F) Alexa Fluor 568 Phalloidin staining showing actin polymerization in AML12 cells treated with CE/CD combined with/without 2 ng/ml TGF-β1 for 72 h. For RT-qPCR, mouse *Ppia* was used as endogenous control. Bars represent mean ± SD (n=3). *p<0.05; **p<0.01, ns = p>0.05. For Western blotting, GAPDH was used as a loading control. Quantification of protein expression was performed using Image J (National Institute of Health, Bethesda, MD). For IF, DRAQ5 was used to stain the nuclei. Scale bars, 25 µm. Images were chosen representatively from 3 independent experiments.

### Cholesterol depletion promotes EMT and actin polymerization in a TGF-β1 dependent manner

In view of the induction of EMT and actin polymerization by CD, we investigated whether these effects depend on TGF-β1. AML12 cells were pre-treated 2 h with 10 µM of the TβR-I kinase inhibitor LY2157299 followed by 14 h-CD treatment. They were then incubated with LY2157299 for 72 h. CD treatment induced delocalization of E-Cadherin from the cell membrane, which was rescued by LY2157299 as shown by E-Cadherin immunofluorescence (Figure 5A). Moreover, the induction of fibronectin by CD was also inhibited by LY257299 (Figure 5B). These results were evident also in the effects of CD on the mRNA expression of EMT markers, which were abolished by LY2157299 (Figure 5C). A similar pattern was observed for the expression of E-Cadherin, as measured by Western blotting (Figure 5D). Furthermore, CD-mediated actin polymerization was reduced to normal level by treatment with the inhibitor (Figure 5E). These results suggest that CD-induced EMT and actin polymerization depend on TGF-β1 signaling.

**Figure 5.**
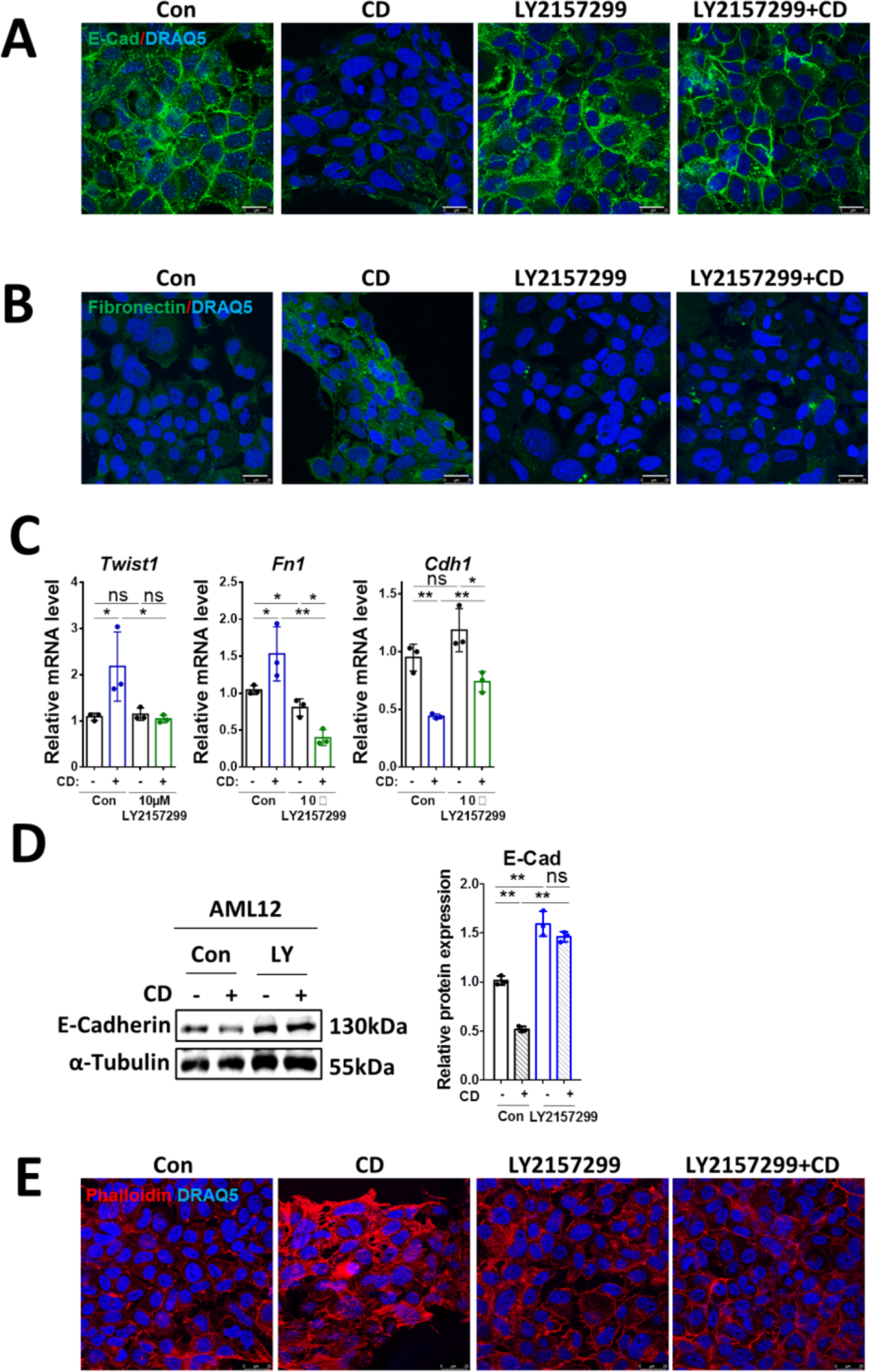
CD-mediated EMT and actin polymerization depend on TGF-β1. (A, B) The expression and location of E-Cadherin and Fibronectin were determined by immunofluorescence in AML12 cells treated with CD combined with/without 10 µM LY2157299 for 72 h. (C, D) Relative mRNA levels of *Twist*, *Fn1* and *Cdh1* (C) and protein levels of E-cadherin (D) were determined by RT-qPCR and Western blotting, respectively, in AML12 cells with the conditioned treatment as shown in the figure. (E) Alexa Fluor 568 Phalloidin staining showing actin polymerization in AML12 cells treated with CD combined with/without 10 µM LY2157299 for 72 h. For RT-qPCR, mouse *Ppia* was used as endogenous control. Bars represent mean ± SD (n=3). \**p*<0.05; \*\**p*<0.01, ns = p>0,05. For Western blotting, GAPDH was used as a loading control. Quantification of protein expression was performed using Image J (National Institute of Health, Bethesda, MD). For IF, DRAQ5 was used to stain the nuclei. Scale bars, 25 µm. Images were chosen representatively from 3 independent experiments.

### Cholesterol depletion induces apoptosis through the caspase 3/7 pathway

To investigate the effect of long-term TGF-β1 treatment in combination with CE or CD on cell apoptosis, we performed an MTT assay. As shown in Figure 6A, this assay displayed that CD treatment significantly inhibited cell proliferation, while CE or TGF-β1 treatment alone did not have an obvious outcome. Next, we conducted apoptosis measurements with the IncuCyte Cytotox Red Dye. Cell numbers analysis confirmed that CD inhibited cell proliferation and markedly promoted cell apoptosis, whereas 72 h incubation with TGF-β1 alone or CE treatment without or with TGF-β1 did not have a significant effect on apoptosis (Figure 6B, C). To further identify the mechanisms underlying CD-induced apoptosis, we performed the Caspase-Glo 3/7 assay and examined expression of cleaved and total caspase3/7 by Western blotting. These studies demonstrated that CD treatment enhanced the signals of cleaved caspase3/7, while treatment with TGF-β1 for 72 h alone did not affect the luminescence measured (Figure 6D). Western blot analysis further supported the notion that CD, but not CE, significantly increased cleaved caspase3/7 (Figure 6E).

**Figure 6.**
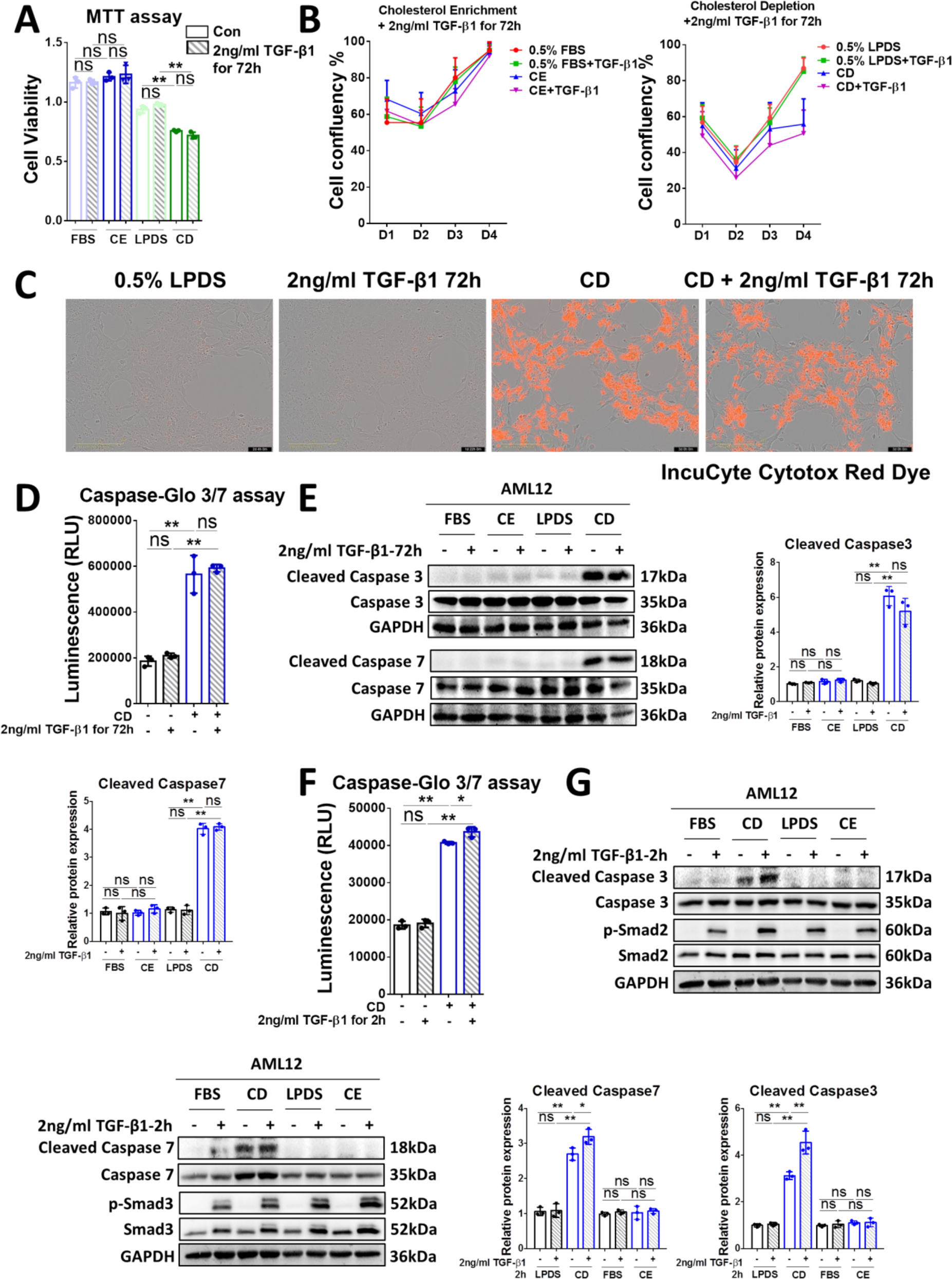
CD induces cell apoptosis through the caspase 3/7 pathway. (A) The viability of AML12 cells treated with CE/CD ± TGF-β1 incubation for 72 h was examined by MTT assay. (B) The confluency of AML12 cells subjected to the conditioned treatments indicated were analyzed by the IncuCyte machine. (C) Cell apoptosis was examined by IncuCyte Cytotox Red Dye in AML12 cells treated with/without CD ± TGF-β1 for 72 h. Scale bars, 50 µm. (D) Cleaved caspase signals were tested by the Caspase-Glo 3/7 assay kit in AML12 cells treated with/without CD ± TGF-β1 for 72 h. (E) Protein levels of cleaved caspase 3/7 and caspase 3/7 were determined by Western blot in AML12 cells treated for CE/CD combined with/without 2 ng/ml TGF-β1 for 72 h. (F) Cleaved caspase signals were tested by the Caspase-Glo 3/7 assay kit in AML12 cells treated with/without CD ± TGF-β1 for 2 h. (G) Protein levels of cleaved caspase 3/7, caspase 3/7, p-Smad2/3, and total Smad2/3 were determined by Western blot analysis in AML12 cells subjected (or not; FBS) to CE/CD, combined with/without 2 ng/ml TGF-β1 for 2 h. For RT-qPCR, mouse *Ppia* was used as endogenous control. Bars represent mean±SD (n=3). \**p*<0.05; \*\**p*<0.01, ns = p>0,05. For Western blotting, GAPDH was used as a loading control. Quantification of protein expression was performed using Image J (National Institute of Health, Bethesda, MD). Images were chosen representatively from 3 independent experiments.

Although it was reported that TGF-β1 signaling can be enhanced by cholesterol depletion ^36^, we did not find obvious TGF-β1-mediated cell apoptosis in the 72 h treated samples. We therefore tested the short-term (2 h) effect of TGF-β1 on cell apoptosis with/without CE or CD treatments. The Caspase-Glo 3/7 assay shows that incubation with TGF-β1 for 2 h alone did not have any effect on the luminescence, while CD treatment alone notably elevated the caspase signals. Of note, CD treatment combined with 2 h of TGF-β1 treatment increased the assays’ readout substantially, as compared to the CD-alone group (Figure 6F). In addition, Western blot reveals that 2 h incubation with TGF-β1 further enhanced cleaved caspase3/7 and p-Smad2/3 levels, compared to the CD treatment group (Figure 6G). In conclusion, CD significantly promoted cell apoptosis, and short-term incubation (2 h) with TGF-β1 further enhanced this effect through the activation of caspase 3/7 pathways.

### Cholesterol depletion promotes cell apoptosis through TGF-β1 signaling

To explore whether the induction of apoptosis by CD involves TGF-β1 signaling, we tested the effect of treating AML12 hepatocytes by LY2157299 (10 μM, 2 h) prior to CD treatment (14 h) followed by incubation up to 72 h with the inhibitor kept in. As shown in Figure 7A, the cell numbers were reduced in the CD-treated group, an effect that was rescued by LY2157299. Quantification using the MTT assay shows that the TβR-I inhibitor promoted, while CD inhibited, cell proliferation compared to the control group (Figure 7B). Notably, LY2157299 retained cell viability despite CD treatment (Figure 7B). The Caspase-Glo 3/7 assay (Figure 7C) and Western blotting for the cleavage of Caspase 3/7 (Figure 7D) further confirmed that blocking TGF-β1 signaling partially inhibited CD-induced expression of cleaved caspase 3 and 7. Taken together, these findings suggest that CD-induced cell apoptosis is dependent on TGF-β1 signaling.

**Figure 7.**
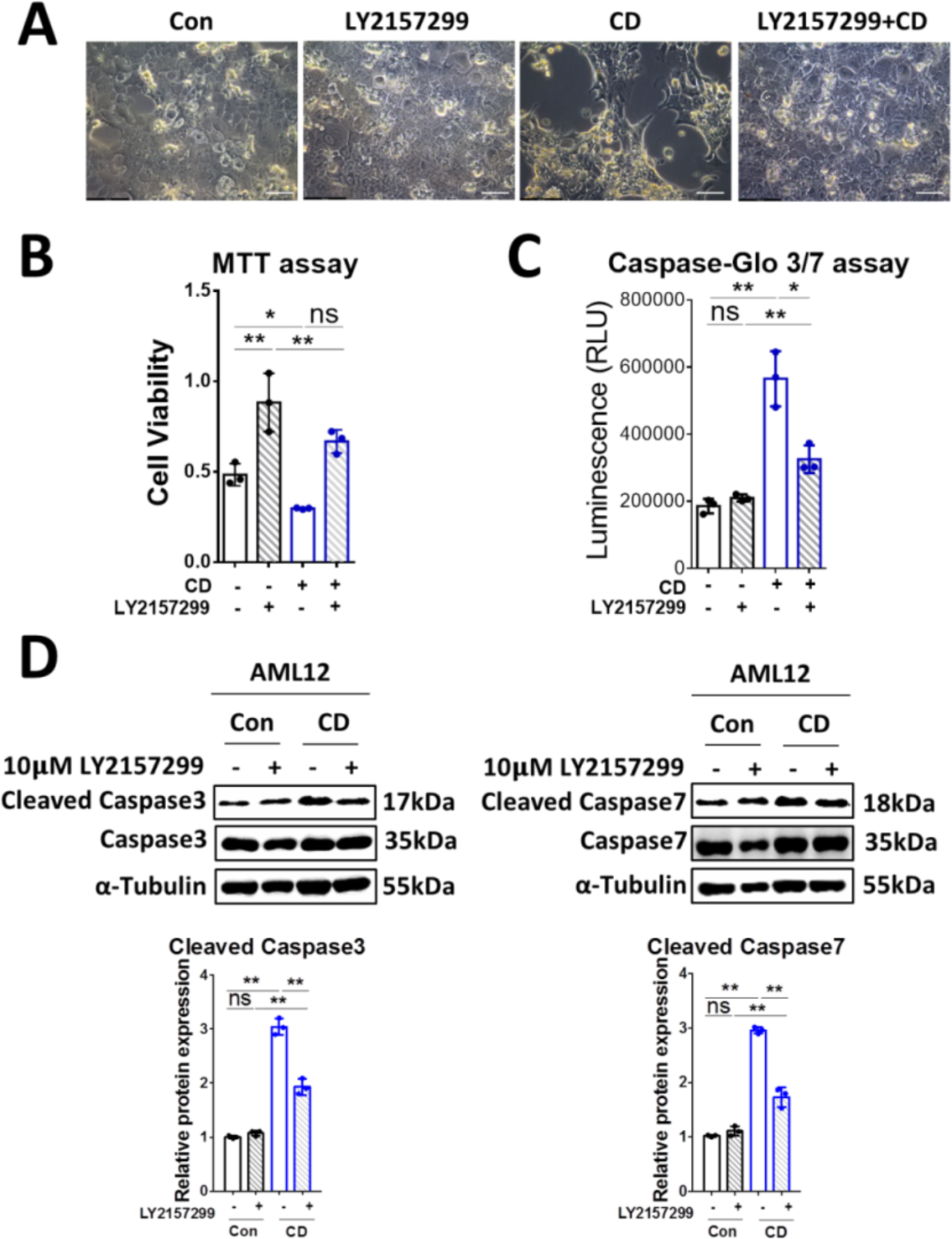
CD promotes cell apoptosis through TGF-β1 signaling. (A) Brightfield images of AML12 cells treated (72 h) with CD alone, LY2157299 (10 μM), or their combination. Untreated cells served as control. Scale bars, 50 µm. (B) The viability of AML12 cells treated for 72 h with/without CD ± 10 µM LY2157299 was examined by MTT assay. (C) Cleaved caspase signals were tested by the Caspase-Glo 3/7 assay kit on AML12 cells subjected (or not; control) to CD treatment ± 10 µM LY2157299 for 72 h. (D) Western blotting of cleaved and total caspase 3/7 in AML12 cells subjected to CD treatment with/without 10 µM LY2157299 for 72 h. For RT-qPCR, mouse *Ppia* was used as endogenous control. Bars represent mean ± SD (n=3). \**p*<0.05; \*\**p*<0.01, ns = p>0,05. For Western blotting, GAPDH was used as a loading control. Quantification of protein expression was performed using Image J (National Institute of Health, Bethesda, MD). Images were chosen representatively from 3 independent experiments.

### Supernatants from CE-treated and CD-treated AML12 cells have opposite effects on HSC activation

Next, we investigated the crosstalk between hepatocytes (AML12) and immortalized hepatic stellate cells (LX-2). To do so, AML12 cells were subjected (or not; control) to the CE or CD treatments, followed by 72 h incubation without the CE or CD-inducing reagents. Supernatants from the AML12 cells were collected, and LX-2 cells were incubated for 72 h with these supernatants. Then, the conditioned LX-2 medium or the cells themselves were collected for further analysis by the SEAP reporter assay to quantify the amount of active TGF-β1 present in conditioned LX-2 supernatants, and the cells were subjected to RT-qPCR of HSC-specific hepatic fibrosis markers. The SEAP assay results indicated that the active TGF-β1 present in conditioned LX-2 supernatants was decreased in the CE group, whilst it was increased in the CD group (Figure 8A). Consistent with the changes observed for levels of active TGF-β1, HSC activation or HSC-specific fibrosis markers, including *COL1A1*, *COL3A1*, *MMP2*, and *MMP9* were notably inhibited in LX-2 cells incubated with supernatants of CE-treated AML12 cells. Conversely, these parameters were significantly increased in the groups of LX-2 cells incubated with supernatants of CD-treated AML12 cells (Figure 8B). Furthermore, we employed an antibody recognizing the latent TGF-β1 latency associated peptide (LAP) breakdown product (LAP-D R58) that stays in the ECM or on the cell membrane upon latent TGF-β activation. Consistent with the SEAP assay result (Figure 8A), immunofluorescent staining displayed an induction of LAP-D R58 in LX-2 cells treated with cholesterol-lowering agents (Figure 8C).

**Figure 8.**
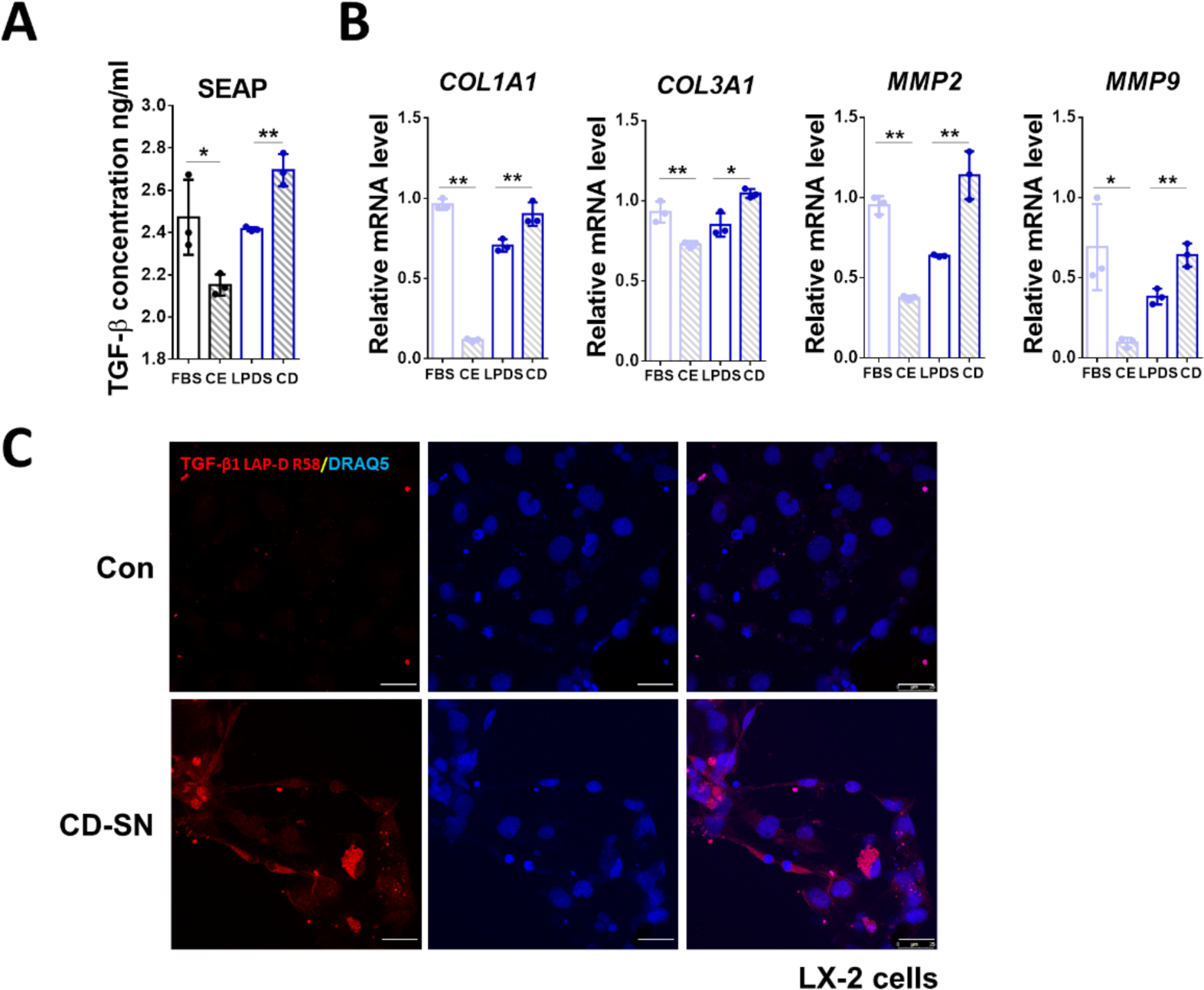
Conditioned media from CE- and CD-treated AML12 cells have opposite effects on HSC activation. (A) Active TGF-β1 concentration in the culture medium of LX2 cells treated with conditioned supernatant of AML12 cells was examined by SEAP activity assay. (B) Relative mRNA levels of *COL1A1*, *COL3A1*, *MMP2*, and *MMP9* were determined by RT-qPCR in LX2 cells treated with conditioned AML12 supernatants as shown in the figure. (C) Expression and location of TGF-β1 LAP-D (R58) were detected by immunofluorescent staining in LX-2 cells incubated with control or CD-treated supernatants of AML12 cells. For RT-qPCR, human *PPIA* was used as the endogenous control. Bars represent mean ± SD (n=3). \**p*<0.05; \*\**p*<0.01, ns = p>0.05. For IF, DRAQ5 was used to stain the nuclei. Scale bars, 25 µm. Images were chosen representatively from 3 independent experiments.

These data suggest that CE-treated hepatocytes protect HSC from activation, whereas CD-treated hepatocytes activate stellate cells and promote subsequent fibrosis.

In conclusion, TGF-β1 inhibits cholesterol biosynthesis and accumulation through downregulation of various genes involved in cholesterol biosynthesis, associated metabolic processes, homeostasis, and transport in hepatocytes, while excess cholesterol levels attenuate TGF-β1 downstream effects in the liver, including EMT, actin polymerization, cell apoptosis, and HSC activation (Figure 9).

**Figure 9.**
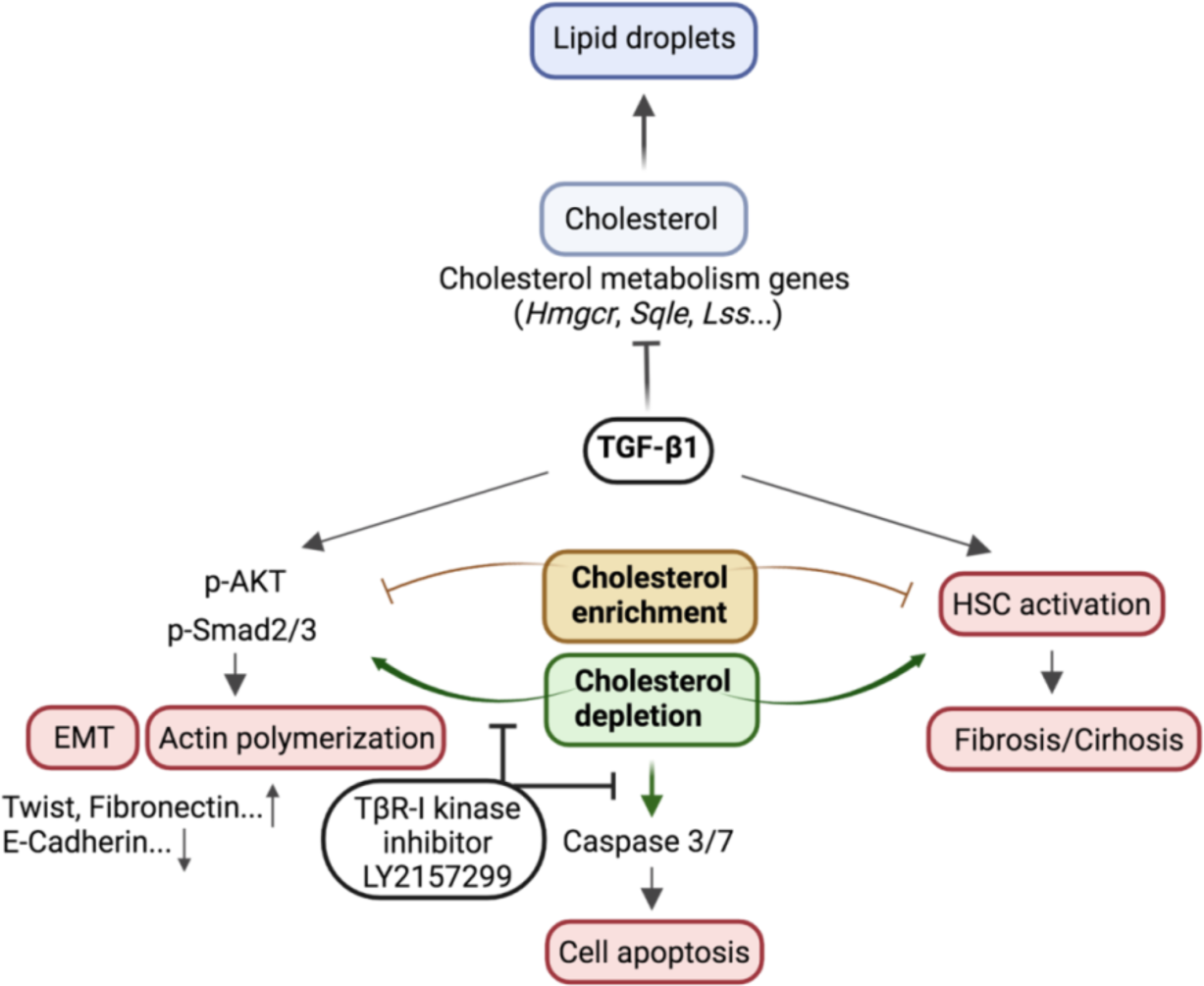
Schematic representation of the crosstalk between the TGF-β1 pathway and cholesterol metabolism. TGF-β1 inhibits cholesterol biosynthesis and lipid droplet accumulation in hepatocytes. CE inhibits TGF-β1-induced EMT, actin polymerization, and HSC activation, while CD enhanced these effects in a TGF-β1-depndent manner. Furthermore, CD induces cell apoptosis through the caspase3/7 pathway reliant on TGF-β1 signaling (Created with BioRender.com).

## Discussion

TGF-β1 signaling and cholesterol level are two significant factors that affect the progression of MASLD ^38^. Yet, knowledge of the effects of TGF-β1 on cholesterol metabolism and the underlying mechanisms is incomplete. In this study, we found that TGF-β1 inhibits expression of multiple genes involved in cholesterol metabolism, including cholesterol homeostasis (*Apoa2*, *Apoa1*, *Hpn*, *Fdft1*, *Angptl3*), cholesterol biosynthesis (*Hmgcs1*, *Sqle*, *Lss*, *Pmvk*, *Cyp5r3*), cholesterol transport (*Abca2*, *Abcg5*, *Ldlr*, *Stard5*), cholesterol esterification (*Soat2*, *Stard4*, *Lcat*) and others. This goes along with reduced accumulation of lipid droplets, as the excess cholesterol is esterified by acyl CoA:cholesterol acyltransferase (ACAT/SOAT) to cholesterol esters, which are either stored as a cholesterol reservoir in cytosolic lipid droplets or released as a major constituent of plasma lipoproteins ^6, 9, 39^, both *in vitro* and *in vivo*. Further research is required to determine how TGF-β1 signaling affects the expression of these genes involved in the cholesterol metabolism. It is possible that some transcription factors of TGF-β1 downstream pathways exhibit common binding sites on the promoters of the aforementioned cholesterol metabolism-associated genes, thus inhibiting their transcription and expression. To test this hypothesis, promoter analysis needs to be performed in the future. Interestingly, we discovered an elevated cholesterol concentration in the plasma of the AAV8-TGF-β1-injected mice (Figure S1), despite the fact that TGF-β1 decreased cholesterol production and accumulation in the liver tissue of the same animals overexpressing TGF-β1. Firstly, the half-life of plasma cholesterol that exceeds several days might serve as an explanation ^40^; secondly, ABCA1 which exports cholesterol into circulation is strongly induced by TGF-β1 treatment ^41^. It also implies that more research is needed on cholesterol import and export in addition to its biosynthesis.

In this study, we also focused on investigating the roles of CE and CD in regulating TGF-β1 downstream biological effects (Figures 4-8). We found that in AML12 cells, CE inhibits TGF-β1-induced EMT and actin polymerization (Figure 4), cell apoptosis (Figure 6), and HSC activation/fibrosis (Figure 8). On the other hand, CD has opposite effects (Figures 4, 6, 8), and its ability to promote these processes depends on TGF-β1 signaling, as indicated by its disruption by a TβR-I inhibitor (Figures 5, 7). These observations may be related to the report that Sirtuin (Sirt) 1/Sirt6 play important roles in TGFβ/Smad pathway inhibition by deacetylation of Smad2 and Smad3 ^42, 43^. Moreover, forkhead box O3 (Foxo3) and krueppel-like factor 6 (Klf6) can bind to the promoters of EMT genes, such as *Snail* and *Twist*, leading to an induction of their transcription ^38^, and cell division cycle 42 (Cdc42) forms complex with N-WASP and Arp2/3 to promote actin polymerization ^44^. The involvement of the above genes in cholesterol homeostasis is suggested by several studies. Thus, increased Sirt1/Sirt6 were shown to decrease cholesterol level ^45^. In addition, it was found that Foxo3/Klf6 and Sirt6 are important for cholesterol homeostasis through controlling SREBP2 expression, resulting in improvement of hypercholesterolemia in diet-induced or genetically obese mice ^46^. These studies imply that there may be a negative feedback regulation between cholesterol to Sirt1/6 and Foxo3/Klf6. As for the small GTPase Cdc42, it was shown to be required for NPC1L1 endocytic recycling compartment, a transporter of cholesterol uptake from intestine and bile, which also requires complex formation with the motor protein myosin Vb and actin filaments ^44, 47–49^. CD could stimulate the formation of GTP-bound, active Cdc42 with increased activity for NPC1L1, resulting in the association of NPC1L1 with myosin Vb and actin, culminating in further contribution to NPC1L1 translocation to the plasma membrane by promoting actin polymerization (Figure 4F). In addition, this is the first report to display caspase 3/7-dependent cell apoptosis induced by CD in hepatocytes; nevertheless, more research is required to understand the underlying mechanisms. We further demonstrate that hepatocytes subjected to CE protect HSCs from activation and then against subsequent fibrosis, whereas CD has the opposite consequences for HSC activation. These results are supported by a study where similar effects were reported to occur following cholesterol enrichment, inducing oxidative stress and HSC mortality that mitigated liver fibrosis ^35^. Further investigation is still needed to fully understand the interaction between CE/CD-treated hepatocytes and HSC activation.

In conclusion, TGF-β1 and cholesterol levels negatively regulate their counterpart, albeit interdependently, in the liver. In terms of therapeutic consequence, we should always account for the adverse effects encountered with cholesterol-lowering treatments when treating MASLD patients with statins and especially TGF-β1-targeted therapies whilst keeping in mind the potential of preventing MASLD progression to HCC.

## Materials and methods

### Reagents and antibodies

Lovastatin (cat. #PHR1285, Sigma-Aldrich, Taufkirchen, Germany); Mevalonate (MVL; cat. #M4667, Sigma-Aldrich); Cholesterol-methyl-β-cyclodextrin (MβCD) soluble complex (cat. #C4951, Sigma-Aldrich); TGF-β1 (cat. #100-21, PeproTech, Hamburg, Germany); LY2157299 (cat. #HY-13226, MedChemExpress; Hoelzel Biotech, Cologne, Germany); Thiazolyl Blue Tetrazolium Blue (MTT) (cat. #11684795910, Sigma-Aldrich); IncuCyte Cytotox Red Dye (cat. #4632, Sartorius, Göttingen, Germany); Caspase 3 antibody (cat. #9662, Cell Signaling Technology, Danvers, MA, USA); cleaved Caspase 3 antibody (cat. #9664, Cell Signaling Technology); Caspase 7 antibody (cat. #9492, Cell Signaling Technology); cleaved Caspase 7 antibody (cat. #8438, Cell Signaling Technology); Phospho-Smad2 (Ser465/467) (cat. #3108, Cell Signaling Technology); Smad2 (cat. #5339, Cell Signaling Technology); Phospho-Smad3 (cat. #ab63403, Abcam); Smad3 (cat. #9523S, Cell

Signaling Technology); Phospho-Akt (Ser473) (cat. #4060, Cell Signaling Technology); Akt (cat. #9272, Cell Signaling Technology); E-Cadherin antibody (cat. #3195, Cell Signaling Technology); Caspase-Glo 3/7 Assay Systems (cat. #8090, Promega, Madison, WI, USA); AlexaFluor 488 AffiniPure F(ab’)_2_ Fragment Donkey Anti-Rabbit IgG (H+L) (cat. #711-546-152, Jackson ImmunoResearch, West Grove, PA, USA); DRAQ5 (cat. #4084L, Cell Signaling Technology); GAPDH antibody (cat. #32233, Santa Cruz Biotechnology, Santa Cruz, CA, USA); Alexa Fluor 568 Phalloidin (cat. #A12380, Invitrogen, Waltham, MA, USA).

### Animal experiments

C57BL/6 mice received a single tail vein injection of 2×10^11^ AAV8-control or AAV8-TGF-β1 for 7 days before collection of liver tissue and blood samples. AAV8-control and AAV8-TGF-β1 were generated by VectorBuilder (Neu-Isenburg, Germany). All experiments were conducted with 8-week-old male mice. Each group contained 3 mice. Animal experiments were carried out in accordance with national guidelines for animal welfare and approved by the local animal care committee of the state of Baden-Württemberg, Germany (Ethics approval 35-9185.81/G-172/15).

### Cell culture and treatment

AML12 murine hepatocyte cells (cat. #CRL-2254) from American Type Culture Collection (ATCC, Manassas, VA, USA) were grown at 37 °C, 5% CO_2_ in high-glucose Dulbecco’s modified Eagle’s medium (DMEM) supplemented with 10% fetal calf serum (FCS), penicillin, streptomycin and L-glutamine as previously described ^50^. They were seeded initially for 12 h either with 2 ml/well of DMEM supplemented with 10% fetal bovine serum (FBS), or with 10% lipoprotein-deficient fetal bovine serum (LPDS) (all from Sigma-Aldrich). For cholesterol depletion (CD), cells were treated with an 50 µM lovastatin and 50 µM MVL in DMEM containing 10% LPDS; the addition of MVL at this concentration prevents excessive reduction of mevalonate, resulting in reduced cholesterol but maintaining normal farnesylation and geranylgeranylation ^51^. For cholesterol enrichment (CE), cells were treated in complete medium (with 10% FBS) with cholesterol-MΠCD complex (5 mM MβCD, 300 µg/ml cholesterol). Following the first 14 h of the above treatments, all groups were starved for 4-6 h with the same aforementioned media respectively, containing either 0.5% FBS or LPDS instead of the original 10% FBS. Following starvation, the cells were stimulated with 2 ng/ml TGF-β1 in DMEM supplemented with either 0.5% FBS or 0.5 % LPDS for 2 or 72 h.

After stimulation, the supernatants were collected for treatment of HSC LX-2 cells (see below). For RNA isolation (for RT-qPCR), the cells were lysed with TRIzol® RNA Isolation Reagents (cat. #15596-018, Thermo Fisher Scientific). For protein analysis, they were lysed with RIPA buffer containing 20 mM Tris–HCl (pH 7.2), 150 mM NaCl, 2% (v/v) NP-40, 0.1% (w/v) SDS, 0.5% (w/v) sodium deoxycholate, Complete™mixture of proteinase inhibitors (Roche, Mannheim, Germany), phosphatase inhibitor cocktail (Sigma-Aldrich) and collected for further protein analysis. For experiments on the effects of the supernatants of the AML12 cells on LX-2 HSCs, the latter were kept in DMEM supplemented with 1% L-glutamine, 1% P/S and 2% FBS and starved for 4 h with starvation medium prior to a 72 h-treatment with AML12 supernatants.

The MFB-F11 cell line is a murine fibroblast cell line isolated from TGF-β-knockout mice (Tgfb-/-) first described by Tesseur et al. ^52^ and were transfected with a Smad-binding element (SBE) controlling a secreted alkaline phosphatase gene (SEAP). MFB-F11 cells were kept in high-glucose DMEM (#11965092, Thermo Fisher Scientific) supplemented with 1% L-glutamine (#25030081, Thermo Fisher Scientific), 1% P/S (#15140122, Thermo Fisher Scientific) and 10% FBS (#10439024, Thermo Fisher Scientific) without sodium pyruvate as an additive. MFB-F11 cells were seeded in 96-well plates (#650160, Greiner Bio-One, Frickenhausen, Germany) at a density of 5 × 10^4^ per well. Following starvation for 2h, conditioned supernatant was added.

All cell lines were routinely checked for absence of mycoplasma contamination via PCR.

### RNA isolation and qRT-PCR

RNA isolation was performed using the TRIzol® reagent (Life Technologies, Carlsbad, CA, USA) according to the manufacturer’s instructions. 500 ng RNA were used for cDNA synthesis using a commercially available cDNA synthesis kit (Thermo Fisher Scientific, Waltham, MA). 20 µl mixtures containing 5 µl cDNA (diluted 1:10), 4 µl Power SYBR® Green Master Mix, 10µM forward and reverse primers were used for real time PCR in a StepOnePlus® Real-Time PCR system (all from Applied Biosystems, Waltham, MA, USA). The RT-qPCR amplification protocol comprised a polymerase activation step for 15 minutes at 95 °C, a subsequent amplification step 15s at 95 °C, 20s at 60 °C, and 20s at 72 °C for 40 cycles. A melting curve was established to validate specificity for each PCR analysis with the following: protocol 15s at 95°C and 1 minute at 60°C, from 60°C to 95°C with +0,3°C every 15 seconds. Human peptidylprolyl isomerase A (hPPIA) was used as a housekeeping gene for normalisation of gene expression. Expressions were calculated with the ΔΔCt method described previously ^53^. Primer sequences were retrieved from the PrimerBank® (Massachusetts General Hospital, Boston, MA, USA) online resource and ordered from Eurofins Genomics (Eurofins Scientific, Luxemburg, Luxemburg) (Table S1).

### Western blotting

Cultured cells were dissolved in RIPA lysis buffer (1% Triton X-100, 50 mMTris [pH 7.5], 300 mM NaCl, 1 mM EGTA, 1 mM EDTA, and 0.1% SDS), supplemented with phosphatase inhibitors (Sigma-Aldrich). A DC® Protein Assay (Bio-Rad, Hercules, CA, USA) was performed according to the manufacturer’s protocol to measure sample protein concentration. Quantification was performed in a Tecan Infinite M200® (Tecan Group AG) microplate reader using the microplate reader’s own protein assay protocol (absorbance at 595nm). Western blot sample protein concentration was calculated using a standard curve of absorbance plotted against pre-defined BSA (bovine serum albumine) concentrations (0, 0.125, 0.25, 0.5, 1, 1.5, 2mg/ml). For each sample, 20 µg of protein were separated by 10% SDS-PAGE gels and were blotted onto a high-resolution nitrocellulose (NC) membranes (MERCK, Darmstadt, Germany). Membranes were blocked with 5% Albumin Bovine Fraction V (SERVA, Heidelberg, Germany) in TBST at room temperature for 1 h. Subsequently, the membrane was incubated with primary antibodies overnight at 4°C. The next day, after washing with TBST for 3x, all membranes were incubated with HRP-linked anti-mouse or anti-rabbit secondary antibodies. Chemiluminescence was determined in a Fusion® SL chemiluminescence reader (Vilber Lourmat Deutschland GmbH, Eberhardzell, Germany) from membranes incubated in Western Lightening® Plus-ECL solution (PerkinElmer, Waltham, MA, USA).

### Determination of cholesterol concentration

Total cholesterol concentration was measured using a cholesterol assay kit (#K603-100 BioVision, Milpitas, CA, USA). 1 × 10^6^ AML12 cells (untreated, or treated to reduce or enrich with cholesterol) were extracted with 200 µl of chloroform:isopropanol:NP40 (7:11:0.1). After centrifugation for 10 min at 15,000 × g, the organic phase was transferred to a new tube, air-dried at 50°C to remove the chloroform, and vacuumed for 30 min to remove trace organic solvent. The dried lipids were dissolved with 200 µl of cholesterol assay buffer and vortexed to homogeneity. 1-50 µl of the extracted sample was used per assay and adjusted to a volume of 50 µl with cholesterol assay buffer. 50 µl of Reaction Mix containing 44 µl cholesterol assay buffer, 2 µl cholesterol probe, 2 µl cholesterol enzyme mix, and 2 µl cholesterol esterase were added to each sample for incubation at 37°C for 1 h. Absorbance was measured at 570 nm using the Infinite 200 Spectrophotometer (Tecan Group AG, Männedorf, Switzerland).

### MTT assay

Proliferation measurements were performed using the thiazolyl blue tetrazolium bromide (MTT) assay. 4,000 cells per well were seeded in quadruplicate in a 96-well plate. After attachment, cells were treated as described in the section *cell culture and treatment*. At the end of all other experiments, the remaining cell culture medium was discarded, and 100 µl of new culture medium and 10 µl MTT reagent was added to each well and incubated for 4 h at 37°C to enable the formation of formazan crystals. After aspiration of the medium, 100 µl/well DMSO were added, and the plate was incubated on a shaker at room temperature for 10 min to dissolve the formazan crystals. Absorbance was measured at 570 nm using the Infinite 200 Spectrophotometer.

### Immunofluorescence staining

AML12 cells were fixed with 4% PFA for 15 min at room temperature, washed three times with PBS, and permeabilized and blocked with 1% BSA and 0.5% Triton X-100 in PBS for one hour. The cells were incubated overnight at 4°C with the E-cadherin or Fibronectin antibody (1:200 in PBS). After washing three times the cells were incubated with the secondary antibody (AlexaFluor 488 AffiniPure F(ab’)_2_ Fragment Donkey Anti-Rabbit IgG (H+L), #711-546-152, 1:200, Jackson ImmunoResearch, West Grove, PA, USA) and the fluorescent DNA dye DRAQ5 (1:1000, #4084L, Cell Signaling Technology) in PBS for one hour at room temperature. After washing three times, the cells were mounted using the DakoCytomation fluorescent mounting medium (#S3023, DAKO, Germany). Stained samples were analysed with TCS SP8 upright Confocal Microscope (Leica, Wetzlar, Germany).

### Phalloidin staining

AML12 cells were fixed, washed and permeabilized as above, and incubated with Alexa Fluor 568 Phalloidin (1:250) and DRAQ5 (1:1000) in PBS for one hour at room temperature. After washing three times, the cells were mounted using Dakocytomation fluorescent mounting medium. Stained samples were analysed by confocal microscopy as above.

### Apoptosis measurement by IncuCyte Cytotox Red Dye

Apoptosis was measured with IncuCyte Cytotox Red Dye. 4,000 AML12 cells were seeded per well in quadruplicate in a 96-well plate. Following attachment, cells were treated as described in *cell culture and treatment*. TGF-β1 was added first, followed by 100 µl of 2×Cytotox Dye to each well containing 100 µl of cell culture medium. Then, the plate was introduced into the IncuCyte chamber placed within a CO_2_ incubator; images were captured every 2 h for 72 h by the IncuCyte Live-Cell analysis system (Leica).

### Caspase-Glo 3/7 assay

Caspase 3/7 signals were measured using the Caspase-Glo 3/7 assay. 4,000 cells per well were seeded in quadruplicate in 96-well plates. Following attachment, cells were treated as described in *cell culture and treatment*. At the end of all other experiments, the cell culture medium was discarded and 100 µl of Caspase-Glo 3/7 reagent were added to each well. Then, the plate was incubated on a shaker at 300-500 rpm for 30 s, followed by incubation at room temperature for 1 h. All samples were then transferred to a white-walled 96-well plate and their luminescence was measured in the Infinite 200 Spectrophotometer.

### SEAP assay

Following the collection of conditioned medium from MFB-F11 TGF-β reporter cells after 48 h of incubation, secreted embryonic alkaline phosphatase (SEAP) activity and TGF-β concentration were determined using the Great EscAPe® SEAP (631738, TaKaRa Bio, Kusatsu, Japan) chemiluminescence kit to quantify SEAP activity (excitation wavelength 360 nm; emission wavelength 449 nm). Chemiluminescence was measured in a Tecan Infinite M200 microplate reader (signal integration time 10000 ms). TGF-β concentration was calculated from SEAP activity based on a standard curve from pre-defined TGF-β concentrations (0, 0.1, 0.25, 0.5, 1, 2, 5, 10, and 20 ng/ml).

### RNA-seq microarray analysis

Total RNA was isolated from AML12 cells or MPHs treated with TGF-β1 (or not; control) for 24 h. Acceptable RNA quality was confirmed by capillary electrophoresis on an Agilent 2100 bioanalyzer (Agilent, Santa Clara, CA, USA). Gene expression profiling was performed using arrays of the mouse MoGene 2.0 type. Biotinylated antisense cRNA was then prepared according to the Affymetrix standard labelling protocol with the GeneChip WT Plus Reagent Kit and the GeneChip Hybridization, Wash and Stain Kit. Afterward, the hybridization on the chip was performed on a GeneChip Hybridization oven 640. Dyeing took place in the GeneChip Fluidics Station 450. Thereafter, chips were scanned with the GeneChip Scanner 3000. All equipment used for RNA-seq microarray analysis was obtained from Affymetrix (Affymetrix, Santa Clara, CA, USA) where not otherwise specified.

### Statistical analysis

Statistical analyses were performed with GraphPad Prism version 6.0 software. The two-tailed Student’s t-test was used to compare two independent groups. One-way ANOVA was adopted to test for statistical differences between the means of two groups. Variables were described by mean and standard deviation (SD). Statistical significance was indicated as follows: \**P* < 0.05; \*\**P* < 0.01, ns > 0.05. Quantification of protein expression and imaging studies was performed using Image J (National Institute of Health, MD).

All authors had access to the study data and have reviewed and approved the final version of this manuscript. Results will be made accessible to fellow researchers upon request to the corresponding author(s).

## Acknowledgment

We acknowledge the support of the LIMa Live Cell Imaging at Microscopy Core Facility Platform Mannheim (CFPM).

## Abbreviations

AAV: adenovirus-associated virus CD: cholesterol depletion CDC42: cell division cycle 42 CE: cholesterol enrichment
ECM: extracellular matrix
EMT: epithelial-Mesenchymal Transition FOXO3: forkhead box O3
HCC: hepatocellular carcinoma
HMGCR: 3-hydroxy-3-methylglutaryl coenzyme A reductase HSC: hepatic stellate cell
KLF6: Krueppel-like factor 6 LPDS: lipoprotein deficient serum LPS: lipopolysaccharide
LSS: lanosterol synthase
MASLD: metabolic dysfunction-associated steatotic liver disease MASH: metabolic dysfunction-associated steatohepatitis
MΠCD: cholesterol-methyl-Π-cyclodextrin complex MVL: mevalonate
PPIA: peptidylprolyl isomerase A
SEAP: secreted embryonic alkaline phosphatase SIRT1/6: Sirtuin 1/6
SQLE: squalene monooxygenase
SREBPs: sterol regulatory element binding proteins
TGF-β1: transforming growth factor-β1
TβR-I/II: transforming growth factor-β receptor-I/II

