## Supplementary document for "TGF-β1 inhibits cholesterol metabolism in hepatocytes to facilitate cell death, EMT and signals for HSC activation"

[1] Department of Medicine II, University Medical Center Mannheim, Medical Faculty Mannheim, Heidelberg University, Mannheim, Germany; [2] Department of Internal Medicine, The Second Hospital of Dalian Medical University, Dalian, China; [3] Clinic of Gastroenterology, Hepatology and Infectious Diseases, Otto-von-Guericke-Unitersity, Magdeburg, Germany; [4] Shmunis School of Biomedicine and Cancer Research, George S. Wise Faculty of Life Sciences, Tel Aviv University, Tel Aviv 6997801, Israel; [5] Department of Neurobiology, George S. Wise Faculty of Life Sciences, Tel Aviv University, Tel Aviv 6997801, Israel; [6] Institute of Molecular Pathobiochemistry, Experimental Gene Therapy and Clinical Chemistry (IFMPEGKC), RWTH Aachen University Hospital, D-52074 Aachen, Germany; [7] Mannheim Institute for Innate Immunoscience (MI3), Medical Faculty Mannheim, Heidelberg University, Mannheim, Germany; [8] Clinical Cooperation Unit Healthy Metabolism, Center of Preventive Medicine and Digital Health, Medical Faculty Mannheim, Heidelberg University, Mannheim, Germany;

**Materials and Methods**

Table S1. Primers for qRT-PCR

| Primer | Forward | Reverse |
|---|---|---|
| hCOL1A1 | GAGGGCCAAGACGAAGACATC | CAGATCACGTCATCGCACAAC |
| hCOL3A1 | TTGAAGGAGGATGTTCCCATCT | ACAGACACATATTTGGCATGGTT |
| hMMP-2 | TACAGGATCATTGGCTACACACC | GGTCACATCGCTCCAGACT |
| hMMP-9 | TGTACCGCTATGGTTACACTCG | GGCAGGGACAGTTGCTTCT |
| hPPIA | AGGGTTCCTGCTTTCACAGA | CAGGACCCGTATGCTTTAGG |
| mCdh1 | CAGGTCTCCTCATGGCTTTGC | CTTCCGAAAAGAAGGCTGTCC |
| mFN1 | GATGTCCGAACAGCTATTTACCA | CCTTGCGACTTCAGCCACT |
| mHmgcr | AGCTTGCCCGAATTGTATGTG | TCTGTTGTGAACCATGTGACTTC |
| mLss | GTGTCTTGGCTGGGTGATAA | GACACCAACACTGACCCTATC |
| mPPIA | GAGCTGTTTGCAGACAAAGTT | CCCTGGCACATGAATCCTGG |
| mSqle | ATAAGAAATGCGGGGATGTCAC | ATATCCGAGAAGGCAGCGAAC |
| mTGF-β1 | AGGGCTACCATGCCAACTTC | CCACGTAGTAGACGATGGGC |
| mTwist1 | CAGCAAGATCCAGACGCTCAAG | ACACGGAGAAGGCGTAGCTGAG |

**Plasma cholesterol assay**

Blood was collected in Li-Heparin vials from the retrobulbar plexus and centrifuged at 13,000 rpm at 4 °C for 6 min. Plasma was subsequently stored at − 80 °C until further analysis. Then cholesterol was measured using a Hitachi automatic analyzer (Core facility-Medical Faculty Mannheim, Germany).

**Statistical analysis**

Statistical analyses were performed with GraphPad Prism version 6.0 software. The two-tailed Student's t-test was used to compare two independent groups. One-way ANOVA was adopted to test for statistical differences between the means of two groups. Variables were described by mean and standard deviation (SD). Statistical significance was indicated as follows: *$P < 0.05$; **$P < 0.01$, ns > 0.05.

**Supplementary figures and figure legends:**

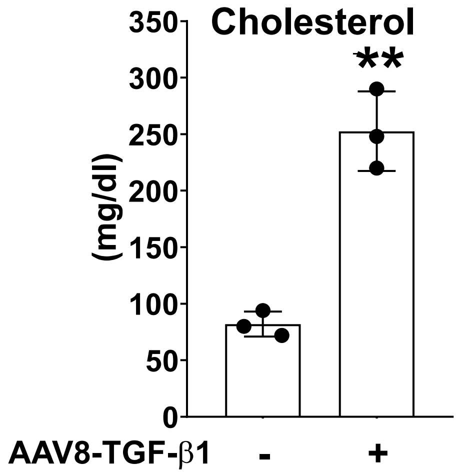


**Figure S1: Overexpression of TGF-β1 induced cholesterol concentration in the serum of mice treated with AAV8-TGF-β1**

Total cholesterol concentration was examined in the serum of WT mice 7 days after the injection of AAV8-Control or AAV8-TGF-β1. Bars represent mean ± SD (n=3). *$p$<0.05; **$p$<0.01, ns > 0.05.
